# The proton-sensing GPR4 receptor regulates paracellular gap formation and permeability of vascular endothelial cells

**DOI:** 10.1101/601088

**Authors:** Elizabeth A. Krewson, Edward J. Sanderlin, Mona A. Marie, Juraj Velcicky, Pius Loetscher, Li V. Yang

**Affiliations:** Department of Anatomy and Cell Biology, East Carolina University, Greenville, North Carolina 27834, USA; Department of Internal Medicine, East Carolina University, Greenville, North Carolina 27834, USA; Novartis Institutes for BioMedical Research, CH-4002 Basel, Switzerland

## Abstract

Tissue acidosis can be a consequence of numerous disease states including stroke, myocardial infarction, limb ischemia, and inflammation. Blood vessels existing in the affected tissues are associated with the progression of acidosis-related diseases. However, the mechanisms by which endothelial cells (ECs) lining the affected blood vessels sense and respond to an acidic microenvironment remain largely unclear. We investigated the functional effects of the proton-sensing G protein-coupled receptor GPR4 in acidosis-induced endothelial inflammation. GPR4 is highly expressed in ECs and known to regulate EC inflammation and endoplasmic reticulum stress responses within acidic microenvironments. Using genetic and pharmacological approaches, we demonstrate that GPR4 activation by acidosis increases EC paracellular gap formation and permeability. We further demonstrate that GPR4-mediated paracellular gap formation is dependent on the Gα_12/13_ signaling pathway. To assess the functional role of GPR4 in the inflammatory response *in vivo*, we utilized an acute hindlimb ischemia-reperfusion mouse model. We demonstrate that both genetic deletion and pharmacological inhibition of GPR4 reduce tissue edema, exudate formation, endothelial adhesion molecule expression, and leukocyte infiltration in the inflamed tissue. Collectively, these data suggest GPR4/Gα_12/13_ signaling mediates acidosis-induced endothelial paracellular gap formation and permeability. This study implicates GPR4 as a candidate therapeutic target for the remediation of inflammation and tissue edema.

## Introduction

The endothelium is a dynamic barrier that can mediate the transvascular movement of fluids and immune cells between the peripheral blood and interstitial tissues. During active inflammation, the production of numerous inflammatory mediators within the inflammatory loci can increase endothelial gap formation and vascular permeability which facilitate leukocyte infiltration and exudate formation in the inflamed tissues [1–3]. Accumulating evidence suggests that microenvironmental factors, such as acidic pH, can stimulate endothelial cell inflammation [4–8]. Inflammatory tissues are characteristically acidic, owing in part to hypoxia and increased glycolytic metabolism of cells and infiltrated leukocytes resulting in heightened proton accumulation. The acidic tissue microenvironment is associated with a wide range of inflammation-associated disease states such as arthritis, inflammatory bowel disease, myocardial infarction, stroke, and limb ischemia. Previous reports note that local pH ranging from 6.0 to 7.0 is common in the microenvironments of inflamed tissues, solid tumors, and ischemic tissues [9–12]. In ischemic disease, one study demonstrated that within 50 minutes of coronary artery occlusion, local tissue extracellular pH decreased from 7.4 to 5.5 in domestic pigs [13]. Furthermore, in the tourniquet-induced rabbit limb ischemia model, local tissue pH decreased rapidly within 1 hour and dropped from 7.30 to 6.36 during the 4 hour course of limb ischemia [14]. In summary, an acidic interstitial pH is an inflammatory microenvironmental factor in many pathological conditions and has been demonstrated to modulate tissue, blood vessel, and immune cell functions [9–12].

Cells can sense extracellular acidification through multiple molecular sensors such as acid-sensing ion channels (ASICs), transient receptor potential (TRP), and proton-sensing G protein-coupled receptors (GPCRs) [7, 11, 15, 16]. GPR4 is a member of the proton-sensing GPCR family, which also includes GPR65 (TDAG8) and GPR68 (OGR1) [7, 11, 16–18]. GPR4 is highly expressed in vascular endothelial cells (ECs) and has been shown to increase the expression of inflammatory cytokines, chemokines, adhesion molecules, and ER stress related genes upon activation by acidic pH in ECs [4, 6, 8]. GPR4 has also been shown to potentiate inflammation *in vivo*. Recent studies found that the genetic deletion of GPR4 in mouse colitis models decreased the expression of endothelial adhesion molecules VCAM-1 and E-Selectin in the intestinal microvasculature which was associated with reduced mucosal leukocyte infiltration and intestinal inflammation [19, 20]. Furthermore, GPR4 was shown to increase tissue injury in a renal ischemia-reperfusion mouse model [21].

Our current study focuses on the GPR4-mediated endothelial paracellular gap formation and vessel permeability in the inflammatory response. Using genetic and pharmacological approaches, we demonstrate that activation of GPR4 by acidosis induces endothelial paracellular gap formation and permeability through the Gα_12/13_ signaling pathway. Furthermore, we demonstrate that the genetic deletion and pharmacological inhibition of GPR4 decrease blood vessel permeability, tissue edema, and leukocyte infiltration in the acute hindlimb ischemia-reperfusion mouse model. Our data suggest that GPR4 has a proinflammatory role in the regulation of the inflammatory response.

## Materials and Methods

### Chemicals and reagents

For endothelial cell monolayer verification and live-cell staining, CellMask Deep Red Plasma Membrane Stain (#C10046) and NucBlue Live ReadyProbes Reagent (#R37605) were purchased from ThermoFisher Scientific (Waltham, MA, USA). The monoclonal antibody for VE-Cadherin (D87F2) was purchased from Cell Signaling Technology (Danvers, MA, USA). Secondary antibody, Rhodamine Red-X goat anti-rabbit (ThermoFisher Scientific) was used for all immunofluorescence experiments. Cellular permeabilization agent Triton X-100 detergent was purchased from BioRad (Hercules, CA, USA). Rhodamine Phalloidin used for F-actin florescence staining was purchased from Cytoskeleton (Denver, CO, USA). For immunofluorescence staining, we used Vectashield Antifade Mounting Medium with DAPI from Vector Laboratories (Burlingame, CA, USA). To evaluate *in vitro* endothelial cell permeability, fluorescein isothiocyanate (FITC)-dextran (average molecular weight 10,000 Daltons) was purchased from Millipore Sigma (St. Louis, MO, USA). The chemicals used for buffering the pH stimulation media include 4-(2-hydroxyethyl)-1-piperazineethanesulfonic acid (HEPES), N-(2-hydroxyethyl)-piperazine-N′-3-propanesulfonic acid (EPPS), 2-(4-morpholino)-ethanesulfonic acid (MES) and were purchased from Sigma-Aldrich (St Louis, MO, USA).

The GPR4 inhibitor, 2-Ethyl-3-(4-((E)-3-(4-isopropyl-piperazin-1-yl)-propenyl)-benzyl)-5,7-dimethyl-3H-imidazo(4,5-b)pyridine (abbreviated as EIDIP), was purchased from Dalton Pharma Services (Toronto, ON, Canada) as previously described [5, 6]. The new generation of GPR4 inhibitor, 2-(4-((2-Ethyl-5,7-dimethylpyrazolo[1,5-a]pyrimidin-3-yl)methyl)phenyl)-5-(piperidin-4-yl)-1,3,4-oxadiazole (GPR4 antagonist 13, also known as NE-52-QQ57), was provided by Novartis Pharmaceuticals (Basel, Switzerland). The vehicle for GPR4 antagonist 13 was a combination of 0.5% methylcellulose purchased from BioConcept AG (Allschwil, Switzerland) and 0.5% Tween-80 purchased from Sigma-Aldrich (St. Louis, MO, USA) and 99% water.

### Cell culture

Primary human umbilical vein endothelial cells (HUVEC), human pulmonary artery endothelial cells (HPAEC), and human lung microvascular endothelial cells (HMVEC-Lung) (Lonza, Walkersville, MD, USA), and primary human colon microvascular endothelial cells (HMVEC-Colon) (Cell Biologics Inc., Chicago, IL, USA) were cultured at 37°C and 5% CO_2_ in a humidified incubator for experimentation. HUVEC and HPAEC were cultured in endothelial cell growth medium 2 (EGM-2), while HMVEC-Lung and HMVEC-Colon were grown in EGM-2-MV medium (Lonza, Walkersville, MD, USA). Cells were used within 8 passages for all experiments.

HUVECs stably expressing MSCV-IRES-GFP (HUVEC/Vector), MSCV-huGPR4-IRES-GFP (HUVEC/GPR4), MSCV-huGPR4 R115A-IRES-GFP (HUVEC/GPR4-R115A), Flink-control shRNA (HUVEC/Control shRNA), or Flink-huGPR4 shRNA (HUVEC/GPR4 shRNA) were developed by retroviral transduction and sorted by fluorescence-activated cell sorting (FACS) as previously described [4–6]. To generate Gα_12/13_ signaling deficient HUVECs, the p115 RGS construct was cloned into the pQCXIP vector as previously described [22, 23]. The p115 RGS-pQCXIP and empty pQCXIP retroviral vectors were stably transduced into the HUVEC/vector and HUVEC/GPR4 cells which were selected with 2µg/mL puromycin for 48 hours.

### Gap formation assay

Endothelial cells were seeded at 1×10^5^ cells/well in a 24-well tissue culture treated plate and were cultured to a confluent monolayer. Intact cellular monolayers were confirmed using a deep red cellular membrane dye (ThermoFisher, #C10046, Waltham, MA, USA) and a live cell nuclear dye (ThermoFisher, #R37605, Waltham, MA, USA). Endothelial cells were then treated with cell culture medium buffered to physiological pH 7.4 or acidic pH 6.4 for defined time points as previously described [4–6]. For assessment of pharmacological inhibition of GPR4 *in vitro*, the GPR4 inhibitor (EIDIP, Dalton Pharma) was added to the pH stimulation media following the GPR4 inhibitor pretreatment for 1 hour in physiological pH 7.4 [5, 6]. Pictures of endothelial cell monolayers were obtained hourly over a 5-hour period using the EVOS Digital Inverted Microscope imaging system. ImageJ software (NIH) was used to quantify gap percentage per field of view (FOV).

### Fluorescent filamentous-actin stress fiber formation assessment

Following the gap formation assay, endothelial cells were washed with DPBS and fixed with 4% paraformaldehyde for 15 minutes. Cells were then permeabilized for 10 minutes with 0.1% Triton X-100 and stained with Rhodamine Phalloidin for 30 minutes (Cytoskeleton Inc., Denver, CO, USA). Cells were then washed in DPBS and mounted with DAPI-containing Vectashield mounting media (Vector Laboratories, H-1200, Burlingame, CA, USA). Cells were imaged using the EVOS Digital Inverted Microscope imaging system.

### Immunofluorescence

The endothelial cell monolayers in 24-well plates were treated with physiological pH 7.4 or acidic pH 6.4 media for 5 hours. Following pH treatments, cells were fixed with 4% paraformaldehyde for 15 minutes, permeabilized with 0.1% Triton X-100 for 5 minutes and blocked with goat serum in phosphate buffered saline with 0.1% Tween-20 for 1 hour. The cells were then incubated with VE-cadherin primary antibody (Cell Signaling Technology, D87F2) in the blocking solution overnight at 4°C. Cells were washed and incubated for 1 hour with Rhodamine Red-X goat anti-rabbit secondary antibody. The wells were then washed five times with PBS and mounted with a glass cover slip using Vectashield with DAPI. The stained cells were viewed and pictures taken with an EVOS Digital Inverted Microscope.

### Cellular permeability assay

Endothelial cell monolayers were cultured to confluency on a Transwell membrane insert with 0.4µm pores (Corning, NY, USA) within a 24-well plate and subsequently treated with physiological pH 7.4 or acidic pH 6.4 media for 5 hours. Following pH treatment, fluorescein isothiocyanate (FITC)-dextran with average molecular weight of 10,000 daltons (Millipore Sigma, St. Louis, MO, USA) diluted into pH stimulation medium was added into the upper chamber of the Transwell insert at a concentration of 1mg/mL. Subsequently, 1 mL of endothelial growth medium was also added to the lower chamber and the Transwell was incubated for 30 minutes at 5% CO_2_ at 37°C. Following the incubation, 100µL of medium was drawn from the lower chamber and the FITC-dextran signal was quantified with a microplate reader at excitation/emission wavelength of 490/520 nm. The degree of cell permeability was determined by comparing the FITC-dextran fluorescent signal from the treatment group to the control group.

### Tourniquet-based acute hindlimb ischemia and reperfusion murine model

All animal experiments were performed on 9-13-week old male and female mice. GPR4 wild-type (WT) or GPR4-deficient (GPR4 KO) mice backcrossed into the C57BL/6 genetic background for 11 generations were used [17]. To assess the inflammatory response post ischemia-reperfusion, the mouse hindlimb ischemia reperfusion injury model was employed using the pneumatic digital tourniquet cuff system (Model DC1.6, Hokanson, Inc., Bellevue, WA, USA). Prior to tourniquet application, a single dose of 0.03 mg/kg Buprenorphine analgesics was administered subcutaneously to each mouse. Mice were then anesthetized with 1% isoflurane inhalation and placed on a temperature-controlled pad followed by the tourniquet application to the left proximal hindlimb under 200 mm Hg regulated pressure for 3hrs as previously described [24]. During the tourniquet-induced hindlimb ischemia procedure, the mouse breathing rate and body temperature was monitored. Mice were injected subcutaneously with 0.5 mL warm saline at the 1.5 hour and 3 hour time points for maintenance of hydration. After the tourniquet was released, the mice were placed into the microisolation cages with soft bedding for a 21 hour hindlimb reperfusion. Hindlimb circumference was measured before the procedure (baseline) and after the ischemia-reperfusion (IR), and the change in hindlimb circumference (value after IR – value at baseline) was used as an indicator to assess tissue edema. For the evaluation of GPR4 antagonist 13 (also known as NE-52-QQ57, Novartis) in the ischemia-reperfusion mouse model, both C57BL/6J and GPR4 WT mice were used for experiments ranging from 9 to 13 weeks old. Mice were administered vehicle control (0.5% methylcellulose/ 0.5% Tween 80/ 99% water) or GPR4 antagonist 13 at 30 mg/kg b.i.d. by oral gavage one day prior to experimental procedure. Mice were given a second dose of vehicle or GPR4 antagonist 13 approximately 2 hours prior to tourniquet cuff procedure. The last dose of either vehicle or GPR4 antagonist 13 was administered 6 hours into the reperfusion phase. The dosage of 30 mg/kg b.i.d. was chosen because of the best efficacy observed in a previous study [25]. All animal experiments were approved by the Institutional Animal Care & Use Committee of East Carolina University and were in accordance with the *Guide for the Care and Use of Laboratory Animals* administered by the Office of Laboratory Animal Welfare, NIH.

### Tissue collection and histological analysis

After the 24 hour hindlimb ischemia-reperfusion procedure, mice were euthanized, and the lower body was dissected for tissue collection. Upon dissection, inflammatory exudate within the interstitial space between upper leg muscle fibrous sheath and skin from the tourniquet-affected hindlimb was weighed and collected. A portion of the hindlimb skin containing inflammatory exudate was removed approximately 20 × 6 mm and fixed in 10% buffered formalin for tissue processing. The remaining exudate on the skin of the mouse was dissected and weighed. Tissues were embedded in paraffin, sectioned at 7µm, and stained with hematoxylin and eosin (H&E) for histopathological analyses. For assessment of neutrophilic infiltration into inflammatory exudate, pictures were taken from H&E tissue sections at five random points with a 20× objective. Leukocytes with the distinct polymorphic nuclear morphology were counted in each field of view using ImageJ software in a blind manner as previously described [19]. Pictures were taken using the Zeiss Axio Imager A1 microscope.

### Immunohistochemistry

Immunohistochemistry was performed on serial sections of 7µm paraffin embedded skin with inflammatory exudate. Tissues were deparaffinized and hydrated to water by a series of declining ethanol concentrations. Antigen retrieval was performed using Tris-EDTA (pH 9.0) with 0.1% Tween 20 to assess endothelial VCAM-1 and E-selectin protein expression using the SuperPicture 3rd Gen Immunohistochemistry detection system (Invitrogen, Waltham, MA). Endogenous peroxidases were quenched and endogenous mouse IgG in blood serum was blocked using the Mouse-on-Mouse blocking reagent (Vector Laboratories, Burlingame, CA). Subsequently, 10% normal goat serum was used to block tissue followed by primary antibody incubation with anti-VCAM-1 (Abcam, ab134047) and E-selectin (Abcam, ab18981, Cambridge, MA) overnight at 4 °C and followed by incubation with secondary antibody. For detection, the Rabbit Vectastain Elite ABC-HRP Kit (Vector Laboratories, Burlingame, CA, USA) was employed according the manufacture’s protocol. The immunohistochemistry staining of tissue with no primary antibody and non-ischemia-reperfusion mice were used for control analysis (data not shown).

To assess mouse plasma IgG protein in the exudate, the mouse Vectastain Elite ABC-HRP Kit was used (Vector Laboratories, Burlingame, CA). Endogenous peroxidase was quenched, blocking serum and biotinylated anti-mouse IgG secondary antibody was used followed by ImmPACT DAB Peroxidase Substrate (Vector Laboratories, Burlingame, CA) to visualize IgG leakiness from plasma. Immunohistochemical staining of tissue sections with no anti-mouse IgG antibody and the use of anti-Rabbit IgG antibody (Vector Laboratories, Burlingame, CA) were used for negative control analysis (data not shown). Pictures were taken with a Zeiss Axio Imager A1 microscope.

### Enzyme-linked immunosorbent assay (ELISA)

ELISA was performed to measure the C-reactive protein (CRP) level in the serum of mice treated with the GPR4 antagonist 13 or vehicle control. Immediately after the mice were euthanized, blood was collected through cardiac puncture and allowed to form clots. After centrifugation, mouse serum on top of the clots was collected and stored in a − 80 °C freezer for further analysis. The CRP concentration in mouse serum was measured using the mouse CRP ELISA kit (Sigma-Aldrich, RAB1121) according to the manufacturer’s protocol.

### Statistical Analysis

GraphPad Prism software was used for statistical analysis. The results were recorded as the mean ± standard error from at least three independent experiments. For three or more groups, ANOVA was used followed by Bonferroni post hoc test. Statistical analyses between two groups were performed using the *t*-test. *P* < 0.05 was considered statistically significant.

## Results

### Acidosis promotes paracellular gap formation in primary endothelial cells

Tissue acidosis commonly exists in inflammatory microenvironments [7, 11, 12, 16, 26]. However, the involvement of acidosis in endothelial cell (EC) gap formation is largely unknown. Four primary vascular ECs were cultured to a confluent monolayer and were treated with either physiological pH 7.4 or acidic pH 6.4 for 5 hours to assess acidosis-induced paracellular gap formation. Under physiological pH 7.4, all ECs maintained a cellular monolayer with no gap formation over the 5 hour time course. However, under acidic pH 6.4 the cellular monolayers were disrupted and paracellular gap formation was observed (Figure 1A). The gap formation of EC monolayers was determined in human umbilical vein endothelial cells (HUVEC), human pulmonary artery ECs (HPAEC), human colon microvascular ECs (HMVEC-Colon), and human lung microvascular ECs (HMVEC-Lung) by calculating the total percent area of gaps in each field of view (Figure 1B-1E). ECs treated with acidic pH 6.4 for 5 hours developed approximately 4-5 percent of gap area relative to the total area (Figure 1B-1E). No gaps, however, could be detected within physiological pH conditions.

**Figure 1.**
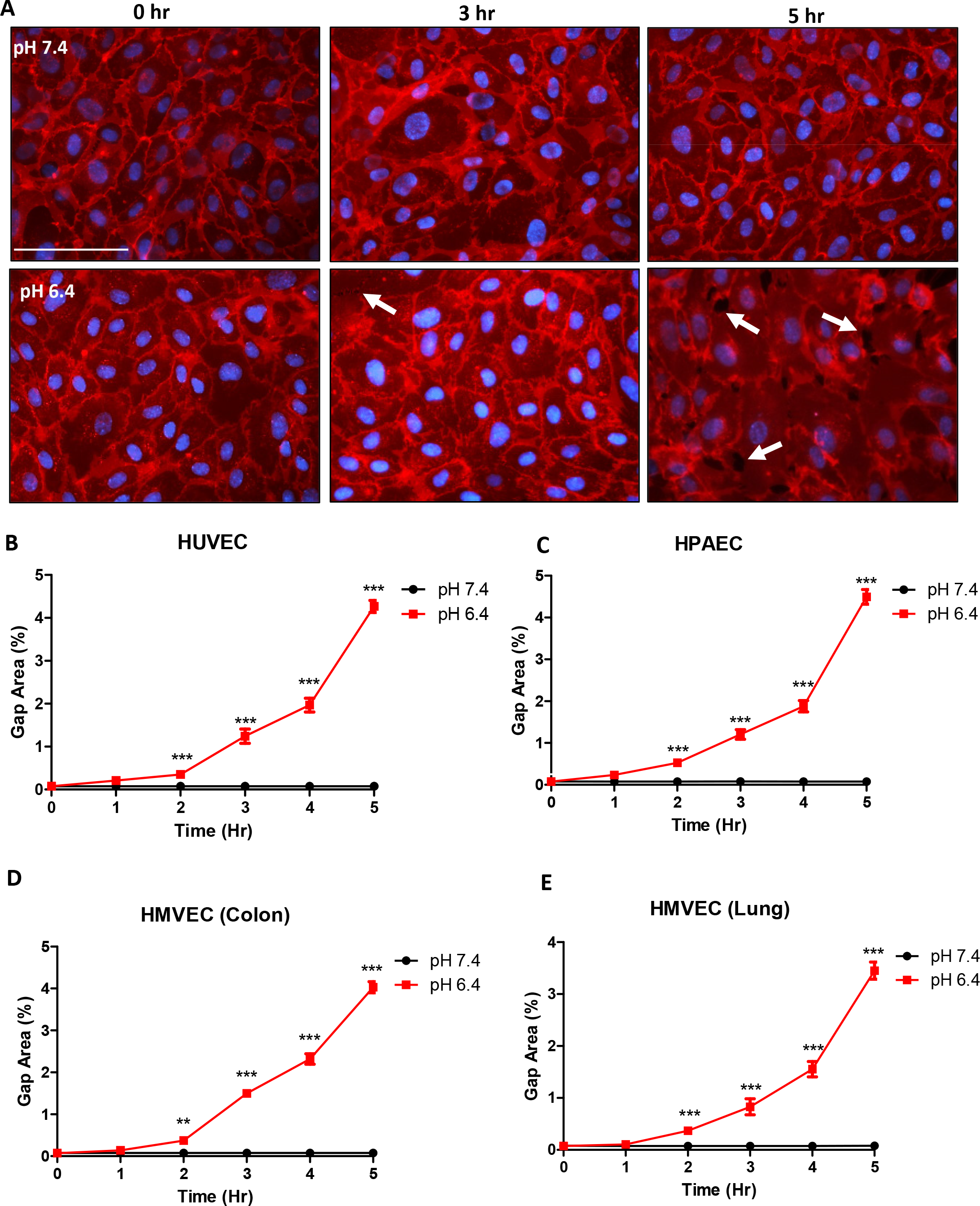
Acidosis stimulates paracellular gap formation in primary vascular endothelial cells (ECs). Plasma membrane staining and paracellular gap area quantitation of ECs treated for up to 5 hours under physiological or acidic pH. Acidosis increases EC gap formation when compared to physiological pH treatment conditions. (A) Representative pictures of plasma membrane staining in Human Umbilical Vein Endothelial Cells (HUVECs) at 0, 3, and 5 hours treated under physiological or acidic pH. Quantitative analysis of gap formation in (B) HUVECs, (C) Human Pulmonary Artery Endothelial Cells (HPAECs), (D) Human Colon Microvascular Endothelial Cells (HMVEC-Colon), and Human Lung Microvascular Endothelial Cells (HMVEC-Lung) over 5 hours. All experiments were performed in triplicate and are representative of four experiments. Data at each time point are presented as mean ± SEM and analyzed for statistical significance between the pH 7.4 group and the pH 6.4 group using the unpaired *t*-test where **p<0.01 and ***p<0.001. White arrows point to paracellular gaps. Scale bar = 100µm.

### Acidosis stimulates endothelial paracellular gap formation through the proton-sensing GPR4 receptor

To determine the role of the pH-sensing receptor GPR4 in acidosis-induced EC gap formation, we used genetic and pharmacological approaches to modulate GPR4 expression and activity, respectively. HUVECs were stably transduced with either control (HUVEC/vector), GPR4 overexpression (HUVEC/GPR4), or GPR4 signaling-defective mutant (HUVEC/GPR4 R115A) overexpression constructs. GPR4 knockdown was achieved with transduction of GPR4 shRNA (HUVEC/GPR4 shRNA) and compared to control shRNA (HUVEC/control shRNA). HUVECs were evaluated under physiological pH 7.4 or acidic pH 6.4 for 5 hours. Treatment with physiological pH 7.4 resulted in no observable gap formation. However, under acidic pH 6.4, HUVEC/vector cell monolayers developed ~4% gap formation (Figure 2A). Overexpression of GPR4 significantly increased acidosis-induced gap formation by ~2.5 fold (~10-11%) under acidic conditions when compared to HUVEC/vector. Conversely, HUVEC/GPR4 R115A mutant decreased the percentage of gap area when compared to HUVEC/vector (~1.8% vs. ~4%, respectively). Furthermore, knockdown of GPR4 by shRNA decreased the percentage of gap area to ~2.5 percent in acidic pH 6.4 conditions when compared to HUVEC/control shRNA (~4.5 percent) (Figure 2B).

**Figure 2.**
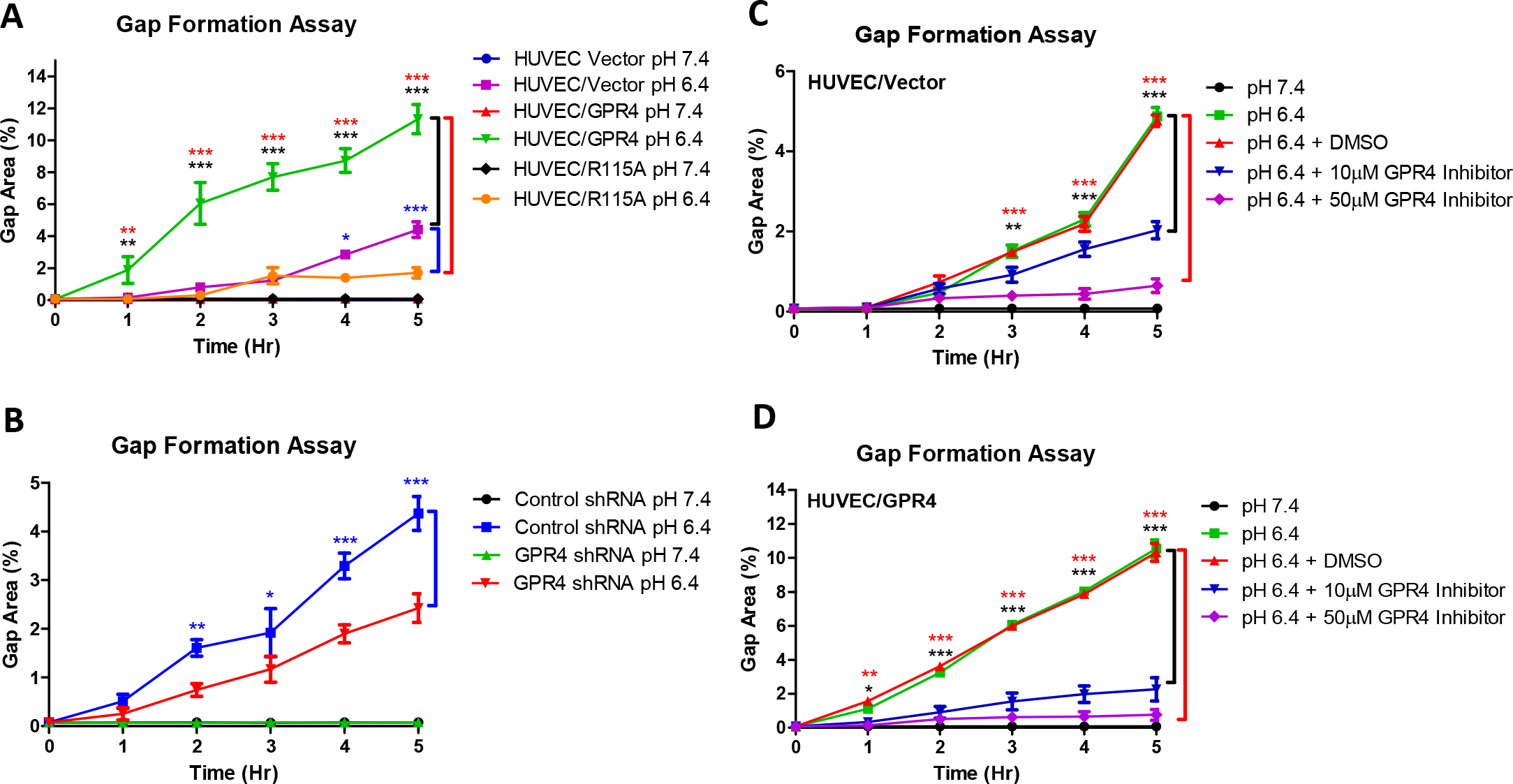
Activation of GPR4 by acidosis induces paracellular gap formation in HUVECs. Quantitative analysis of gap formation utilizing genetic and pharmacological approaches to modulate GPR4 expression or activity. (A) HUVEC/vector, HUVEC/GPR4, and HUVEC/R115A cells. (B) HUVEC/control shRNA and HUVEC/GPR4 shRNA cells. (C) HUVEC/vector and (D) HUVEC/GPR4 cells treated with GPR4 inhibitor or vehicle. All cells were treated with physiological pH 7.4 or acidic pH 6.4 for 0, 1, 2, 3, 4, and 5 hours and the percent of gap area was quantified. DMSO used as vehicle control. All experiments were performed in duplicate or triplicate and are representative three experiments. Data at each time point are presented as mean ± SEM and analyzed for statistical significance using the ANOVA where *p<0.05, **p<0.01 and ***p<0.001. Black, blue, and red symbols indicate statistical analysis between groups indicated by bracket markers.

We next assessed the effects of a GPR4 inhibitor (EIDIP) on acidosis-induced paracellular gap formation in HUVEC/vector and HUVEC/GPR4 cells (Figure 2C and 2D). Pharmacological inhibition of GPR4 attenuated acidosis-induced gap formation. A dose-dependent decrease in the acidosis-induced gap development could be observed with increasing inhibitor concentrations during HUVEC/vector and HUVEC/GPR4 treatments when compared to vehicle (Figure 2C and 2D). The results indicate that acidosis-induced EC paracellular gap formation is dependent on GPR4 activation by acidic pH.

### GPR4-mediated Gα_12/13_ signaling is involved in endothelial paracellular gap formation in response to acidosis

Previous studies demonstrate that the Gα_12/13_/Rho GTPase pathway can regulate cytoskeletal dynamics, endothelial gap formation, and endothelial permeability [27, 28]. In line with these observations, GPR4 can couple to Gα_12/13_ and Rho GTPase when expressed in cancer cell lines [29, 30]. For this reason, we investigated the role of the GPR4/Gα_12/13_ pathway in acidosis-induced EC gap formation. The p115 RGS Gα_12/13_ inhibitory construct [22, 23] was stably transduced into HUVEC/vector and HUVEC/GPR4 cells. We next performed the gap formation assay under physiological and acidic pH conditions for 5 hours. HUVEC/p115 RGS cells treated with acidic pH had significantly reduced gap formation when compared to the vector control (Figure 3A and 3B). Furthermore, thiazovivin (TA) and staurosporine (STA), two chemical inhibitors for Gα_12/13_ downstream effectors Rho-associated kinase (ROCK) and myosin-light chain kinase (MLCK), respectively [30], were used in HUVEC/vector and HUVEC/GPR4 cells under physiological and acidic conditions. Thiazovivin significantly decreased gap formation percentage by ~5 fold in HUVEC/vector and by ~10 fold in HUVEC/GPR4 cells when compared to the vehicle controls under acidic pH. Staurosporine nearly abolished acidosis-induced gap formation in HUVEC/vector and HUVEC/GPR4 cells (Figure 3C and 3D). Collectively, the results suggest that acidosis-induced EC gap formation relies, at least in part, on the GPR4/Gα_12/13_ pathway. Additionally, our results demonstrated that acidosis also induced F-actin stress fiber formation and decreased VE-cadherin expression at the site of paracellular gaps in ECs (Supplemental Figures 1 and 2).

**Figure 3.**
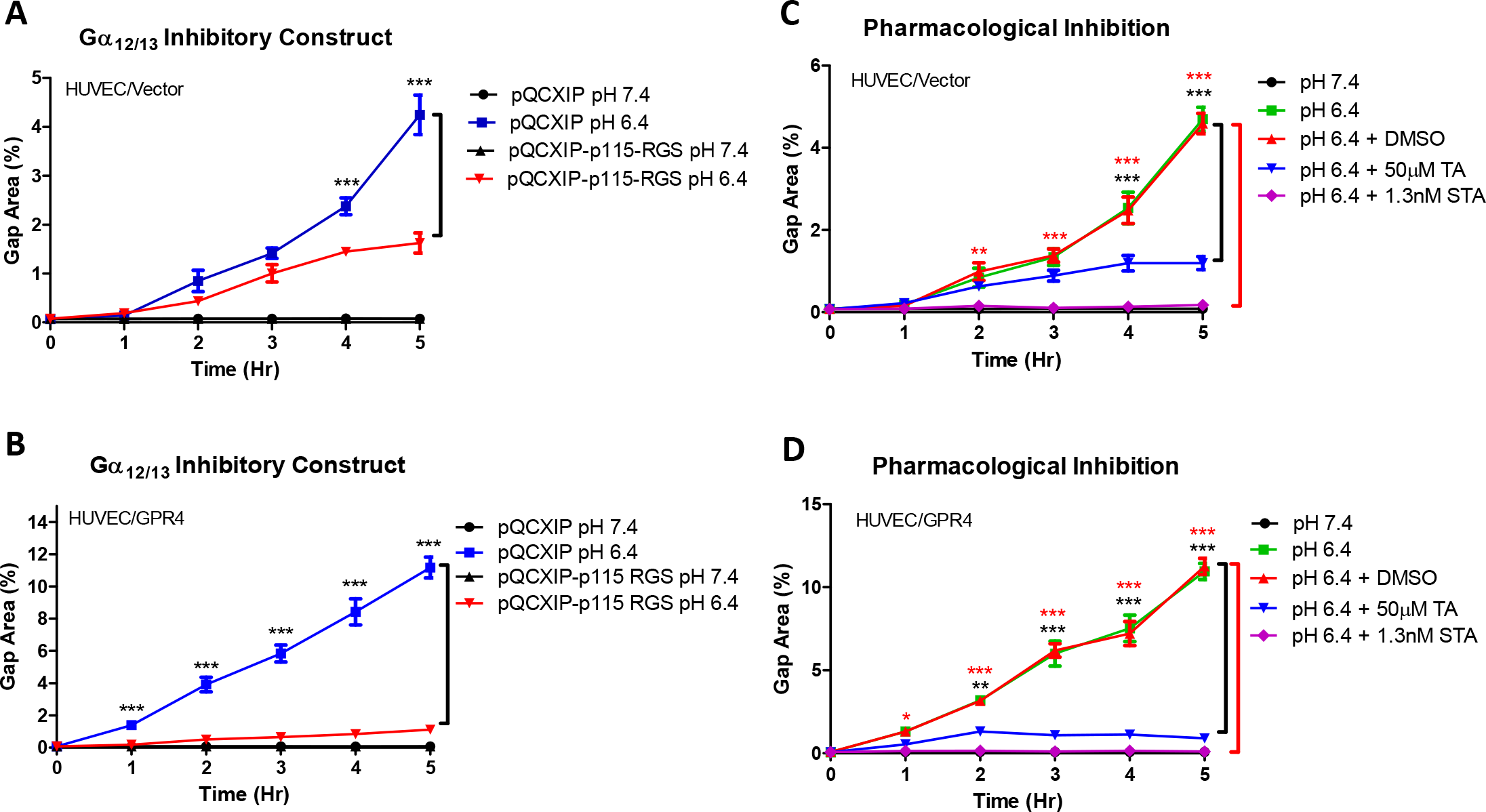
The Gα12/13 pathway is involved in GPR4-mediated paracellular gap formation in HUVECs. Quantitative analysis of GPR4/Gα12/13-mediated paracellular gap formation in HUVECs. GPR4 mediates paracellular gap formation is dependent, at least in part, through Gα12/13 in response to acidic pH. The Gα12/13 p115-RGS inhibitory construct in (A) HUVEC/vector and (B) HUVEC/GPR4 cells. Pharmacological inhibition of the Gα12/13 pathway using 50µM thiazovivin (TA) or 1.3nM staurosporine (STA) in (C) HUVEC/vector or (D) HUVEC/GPR4 cells. HUVECs were treated with either physiological pH 7.4 or acidic pH 6.4 for up to 5 hours. All experiments were performed in duplicate and are representative of at least three experiments. Data at each time point are presented as mean ± SEM and analyzed for statistical significance using the ANOVA where *p<0.05, **p<0.01 and ***p<0.001. Black and red symbols indicate statistical analysis between groups indicated by bracket markers.

### Acidosis increases endothelial cell permeability through GPR4

We next assessed if GPR4-dependent paracellular gap formation can functionally result in increased endothelial permeability. Endothelial permeability was assessed by the fluorescein isothiocyanate conjugated-dextran (FITC-dextran) permeability assay whereby diffusion of FITC-dextran through the endothelial monolayer from the upper to lower chamber of the Transwell insert was assessed. Cells were treated with physiological pH 7.4 or acidic pH 6.4 for 5 hours followed by the addition of FITC-dextran. Acidic pH treatment significantly increased FITC-dextran permeability in HUVEC/vector cells when compared to physiological pH. Moreover, when GPR4 is overexpressed and treated with acidic pH there was a further increase in acidosis-induced FITC-dextran when compared to HUVEC/vector cells. When GPR4 expression is knocked down or signaling is defective (R115A mutant) [4], FITC-dextran permeability was significantly decreased compared to controls under acidic conditions (Figure 4). These results suggest that acidosis increases EC permeability through GPR4.

**Figure 4.**
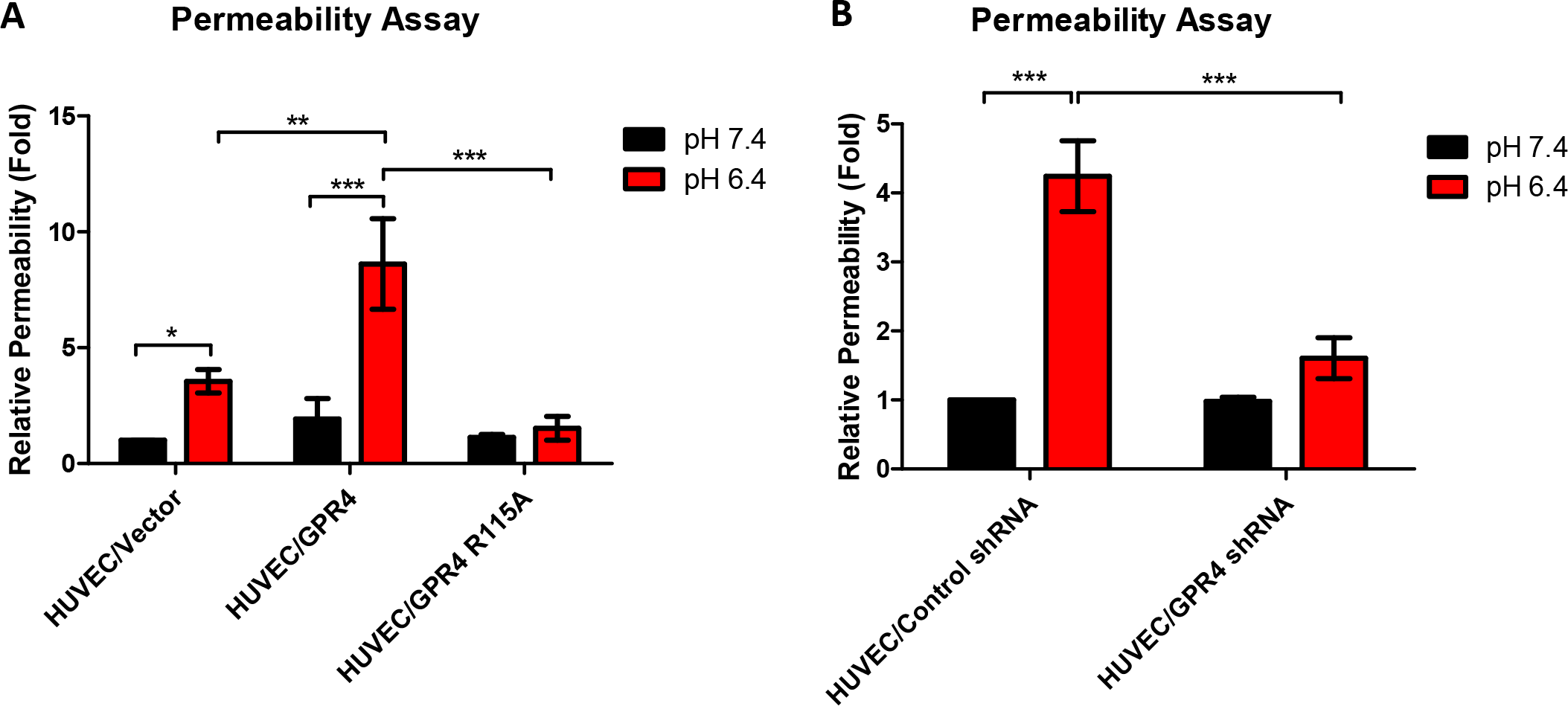
GPR4 activation by acidosis increases cellular permeability in HUVECs. Quantitative analysis of cellular permeability *in vitro*. GPR4 activation by acidic pH increases cellular permeability using the FITC-dextran cellular permeability assay with HUVECs. Relative fold increase of cellular permeability in (A) HUVEC/vector, HUVEC/GPR4, HUVEC/R115A cells, and (B) HUVEC/control shRNA and HUVEC/GPR4 shRNA cells. Cells were treated for 5 hours with either physiological pH 7.4 or acidic pH 6.4. All experiments were performed in duplicate and are representative of at least three experiments. Data are presented as mean ± SEM and analyzed for statistical significance using the ANOVA where *p<0.05, **p<0.01, and ***p<0.001.

### Genetic deletion of GPR4 reduces tissue edema and inflammation in the acute hindlimb ischemia-reperfusion mouse model

Next, we assessed the functional role of GPR4 in a tourniquet cuff–based acute hindlimb ischemia-reperfusion mouse model [24]. This mouse model can cause severe inflammation resulting in increased vessel permeability, tissue edema, and leukocyte infiltration in the affected tissue. The ischemia-reperfusion associated inflammation was induced in wild-type (WT) and GPR4 knockout (KO) mice. The sham and cuff limbs were measured for circumference differences between pre- and post-procedure measurements. GPR4 KO mice had less observable tissue edema in the tourniquet-affected limb following the ischemia and reperfusion event when compared to WT mice (Figure 5).

**Figure 5.**
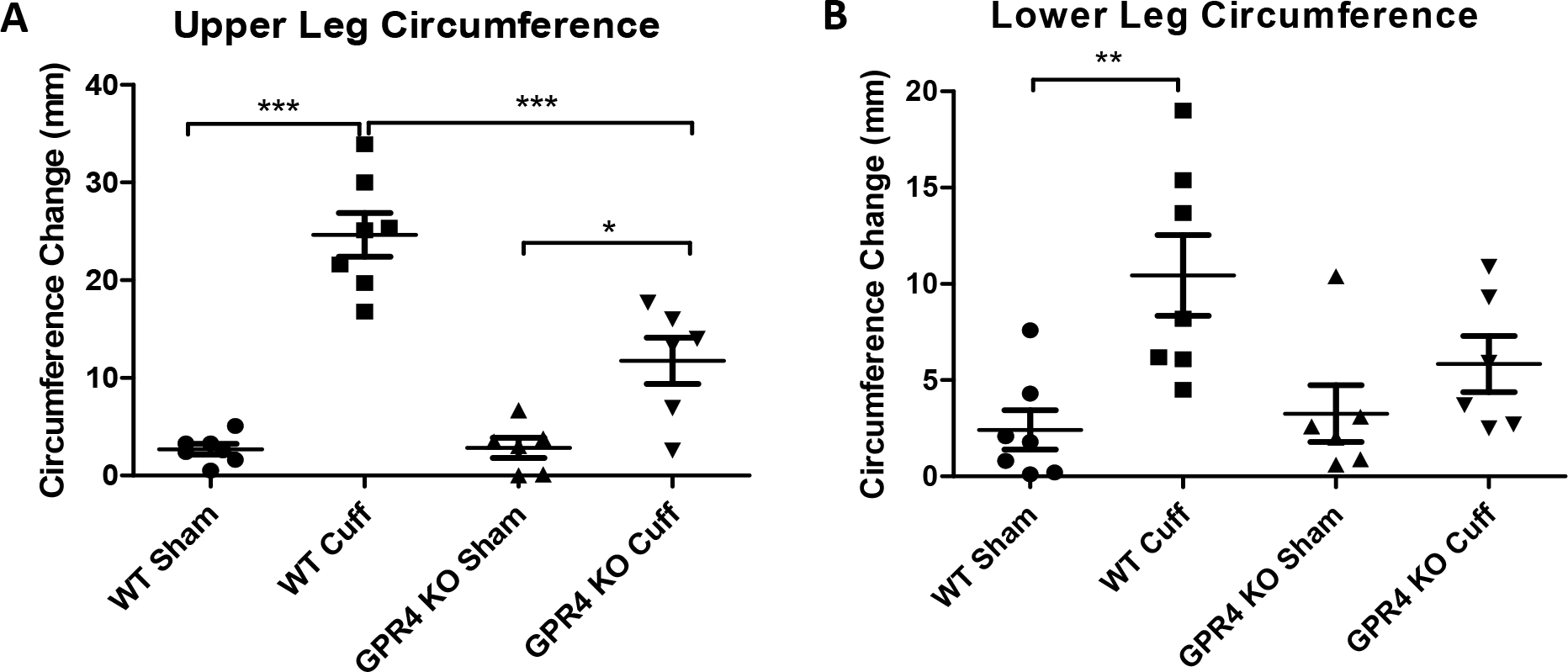
GPR4 deficiency reduces tissue edema in the inflammatory hindlimb ischemia-reperfusion (IR) mouse model. Upper and lower leg circumferences in wild-type (WT, N=7) and GPR4 knockout (GPR4 KO, N=6) mice were measured before and after IR. The change of leg circumference was calculated (value after IR – value before IR) and used as an indicator for tissue edema. GPR4 deficiency reduces the change of upper and lower leg circumference when compared to WT mice in the IR mouse model. Quantitative analysis of (A) upper and (B) lower leg circumference changes. Data are presented as mean ± SEM and analyzed for statistical significance using the ANOVA where *p<0.05 **p<0.01, and ***p<0.001.

Inflammatory exudates in the interstitial space between the skin and the limb muscle/body peritoneum on the tourniquet subjected side were collected and measured followed by histology (Figure 6). GPR4 KO mice had reduced inflammatory exudate when compared to the WT mice (~6mg vs. ~65mg, respectively) (Figure 6A, 6B, and 6I). Furthermore, histological analysis of the inflammatory exudates revealed GPR4 KO mice had reduced leukocyte infiltrates when compared to WT (Figure 6D, 6F, and 6J). No red blood cells were observed in the exudate suggesting the exudate is due to increased vascular permeability but not vessel hemorrhaging (Figure 6A-6F). To further assess the GPR4-mediated vessel permeability, we performed immunohistochemistry for plasma protein immunoglobulin G (IgG) and detected IgG within inflammatory exudates (Figure 6G and 6H), indicating an increased vascular permeability to plasma protein.

**Figure 6.**
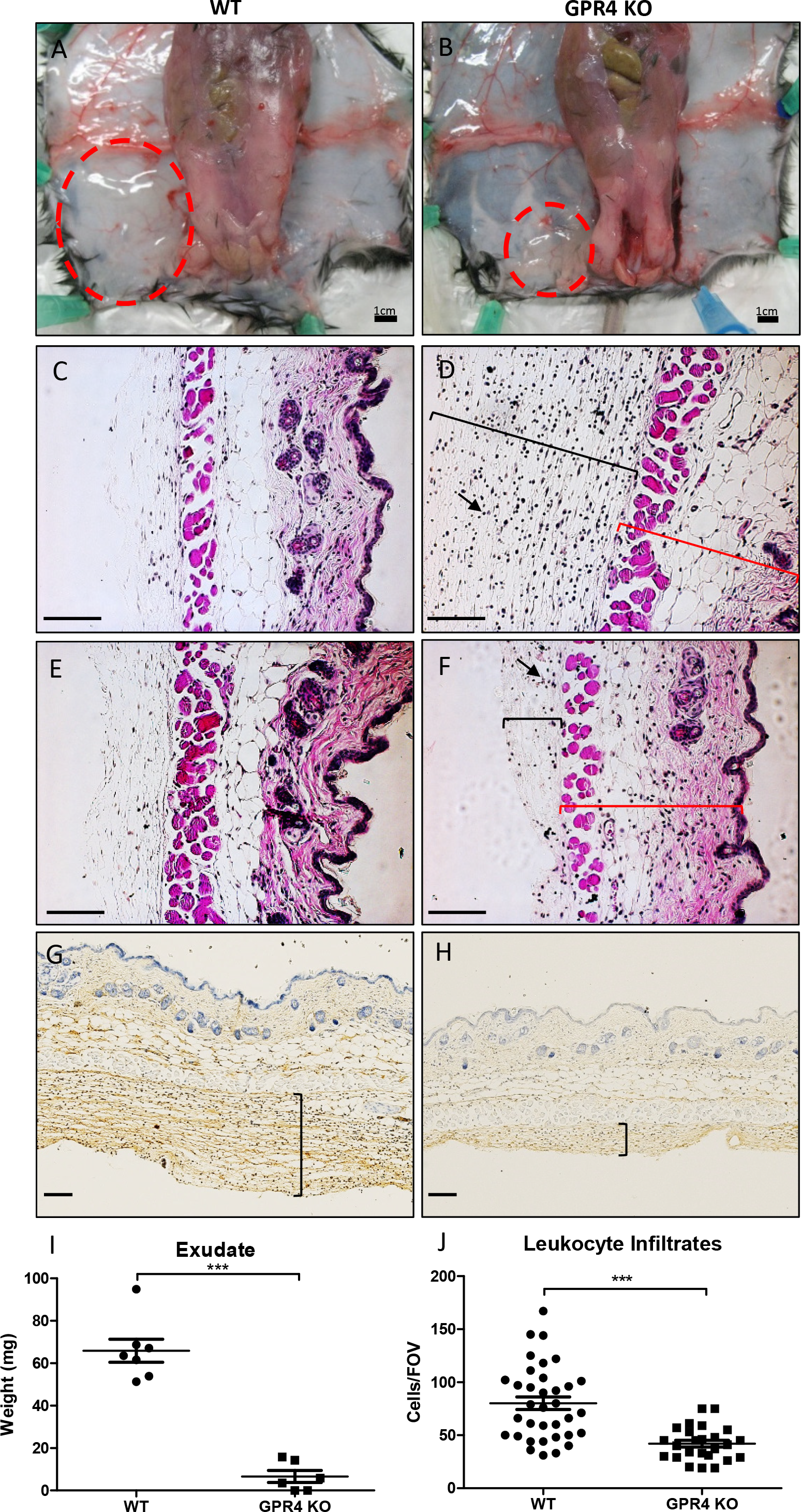
GPR4 deficiency reduces inflammatory exudate production and leukocyte infiltration in the ischemic hindlimb tissues. Inflammatory exudate measurements and leukocyte infiltration quantification in WT and GPR4 KO mice following the ischemia-reperfusion of the hindlimb. Genetic deletion of GPR4 reduces inflammatory exudate production and leukocyte infiltration into the inflammatory exudate. Gross representative images of observable exudate formation in (A) WT and (B) GPR4 KO mice in the interstitial space between the skin and muscle of the ischemic hindlimb. Representative images of H&E staining of (C) WT sham, (D) WT cuff, (E) GPR4 KO sham, and (F) GPR4 KO cuff inflammatory exudate sections. Black brackets indicate exudate distribution. Arrows indicate infiltrated leukocytes in the inflammatory exudate. Red brackets indicate skin tissues. Representative images of mouse plasma IgG protein in (G) WT cuff and (H) GPR4 KO cuff sections. IgG protein can be visualized as brown signal. The plasma protein IgG is used as a marker to indicate blood vessel permeability. Quantitative analysis of (I) exudate weight and (J) leukocyte infiltration from multiple fields of view (FOV). 10× and 20× microscope objectives used. Scale bar indicates 100µm. Data analyzed from 7 WT and 6 GPR4 KO mice. Data are presented as mean ± SEM and analyzed for statistical significance using the unpaired two-tailed *t*-test where ***p<0.001.

Furthermore, less plasma IgG could be observed in the GPR4 KO cuff-affected tissues compared to WT cuff-affected tissues indicating reduced endothelial cell permeability in the KO (Figure 6G and 6H).

To provide a molecular explanation for GPR4 mediated inflammation within vascular endothelial cells, we performed immunohistochemistry to examine the expression of VCAM-1 and E-selectin within the endothelium of the cuff and sham affected hypodermis tissues. Overall, VCAM-1 and E-selectin protein expression was increased within the cuff-affected limb vasculature when compared to sham (Figure 7). Immunohistochemical analysis of VCAM-1 revealed expression on a variety of cell types, such as skeletal muscle, fibroblast, and vascular endothelial cells which is consistent with previous literature [31, 32]. Interestingly, we observed a decrease of VCAM-1 protein expression on vascular endothelial cells in GPR4 KO cuff-affected hypodermis of the limbs when compared to WT (Figures 7A-7D). E-selectin expression was observed on endothelial cells and fibroblasts as previously reported [33–35]. Similar as VCAM-1 expression patterns, there is an observable decrease in E-selectin protein expression in GPR4 KO cuff-affected hypodermis of the limbs when compared to WT (Figures 7E-7H).

**Figure 7.**
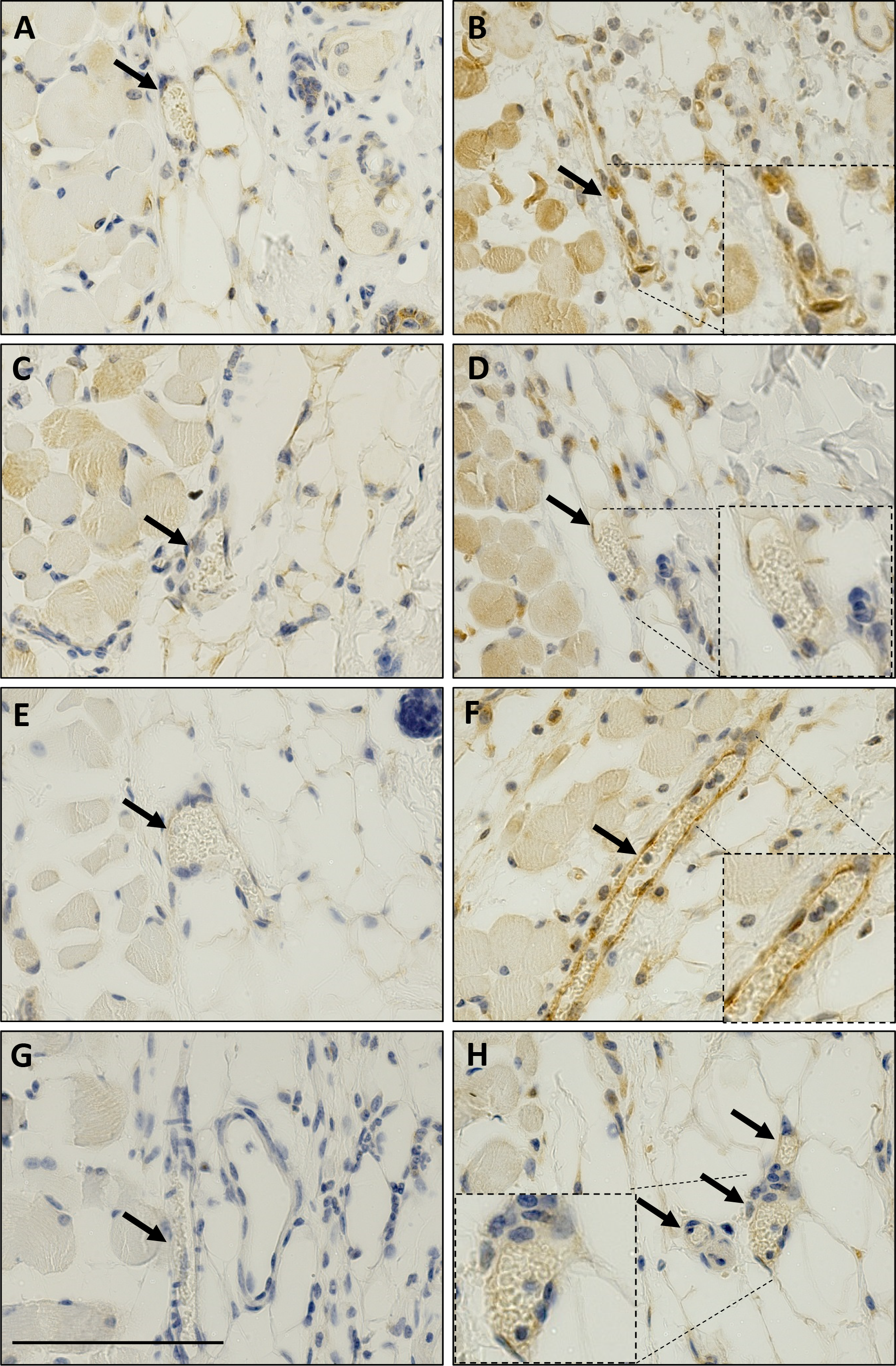
Immunohistochemistry of VCAM-1 and E-selectin protein expression in the loose connective dermal tissue of WT and GPR4 KO mice. Immunohistochemical analysis of adhesion molecules VCAM-1 and E-selectin protein expression in vascular endothelial cells within mouse dermal tissue sections of sham and cuff affected hindlimbs. GPR4-deficient mice had reduced expression of VCAM-1 and E-selectin in vascular endothelial cells in cuff-affected tissues when compared to WT. Representative pictures of VCAM-1 signal in (A) WT sham, (B) WT cuff, (C) GPR4 KO sham, and (D) GPR4 KO cuff tissues. Representative pictures of E-selectin expression can be visualized in (E) WT sham, (F) WT cuff, (G) GPR4 KO sham, and (H) GPR4 KO cuff tissues. Pictures taken with 40× objective. Scale bar indicates 100µm. Arrows indicate blood vessels.

### Pharmacological inhibition of GPR4 reduces tissue edema, leukocyte infiltration, and vascular permeability in the hindlimb ischemia-reperfusion mouse model

To further assess the role of GPR4, we incorporated the use of a highly potent and selective GPR4 antagonist (referred to as GPR4 antagonist 13) [25] within the acute hindlimb ischemia-reperfusion mouse model. Our results demonstrated that the GPR4 antagonist 13 significantly decreased upper and lower limb tissue edema measured by the circumference of the leg compared to limbs of the vehicle control (Figure 8). In addition, there were significantly less leukocyte infiltration within the inflammatory exudate in the GPR4 antagonist 13 treated mice when compared to vehicle treated mice (Figure 9A, 9B, and 9F). Furthermore, the exudate weight was also decreased in GPR4 antagonist 13 treated mice when compared to vehicle (Figure 9E). Additionally, vascular permeability was assessed and there was a reduction in plasma IgG diffusion in the exudate by the treatment of the GPR4 antagonist 13 when compared to vehicle treatment (Figure 9C and 9D). GPR4 antagonist 13 treatment also reduced the level of an inflammatory marker, C-reactive protein (CRP), in the mouse serum (Supplemental Figure 3). To further examine the role of GPR4 within ECs, we performed immunohistochemistry to analyze the expression of VCAM-1 and E-selectin within the endothelium of the cuff affected hypodermis tissues. We observed that ECs in the GPR4 antagonist 13 treated mice had reduced VCAM-1 and E-selectin protein expression compared to the vehicle treated mice (Figure 10). Taken together, these data suggest that GPR4 can mediate inflammation, vessel permeability, and leukocyte infiltration into inflamed tissues.

**Figure 8.**
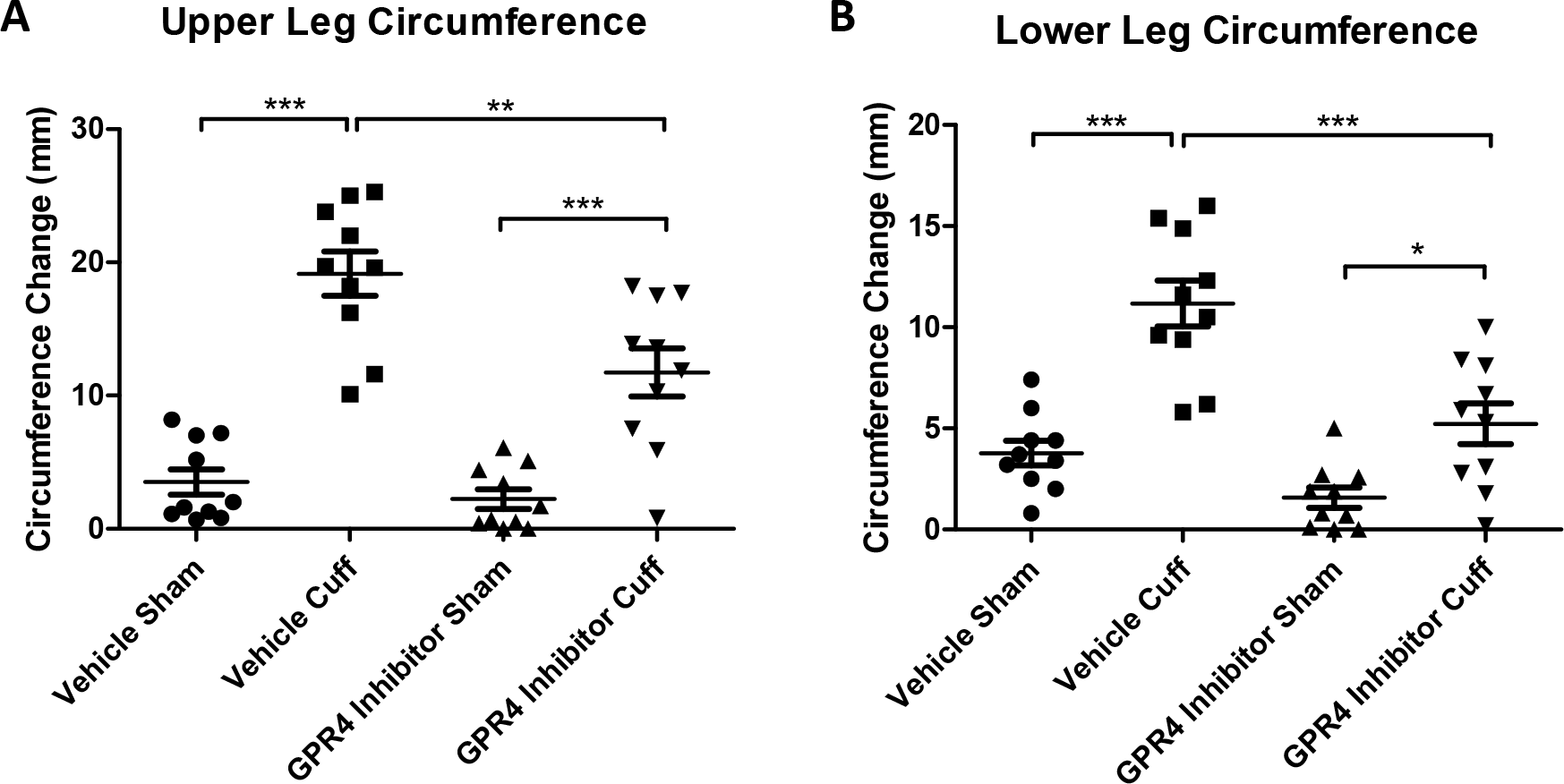
Pharmacological inhibition of GPR4 reduces tissue edema in the inflammatory hindlimb ischemia-reperfusion (IR) mouse model. Upper and lower leg circumference changes following IR in wild-type (WT) mice treated with either vehicle or GPR4 antagonist 13. GPR4 antagonist 13 reduces the changes of upper and lower leg circumference when compared to vehicle control in the IR mouse model. Quantitative analysis of (A) upper and (B) lower leg circumference changes, respectively. N=10 vehicle and N=10 GPR4 antagonist 13 treated mice. Data are presented as mean ± SEM and was analyzed for statistical significance using the ANOVA where *p<0.05 **p<0.01, and ***p<0.001.

**Figure 9.**
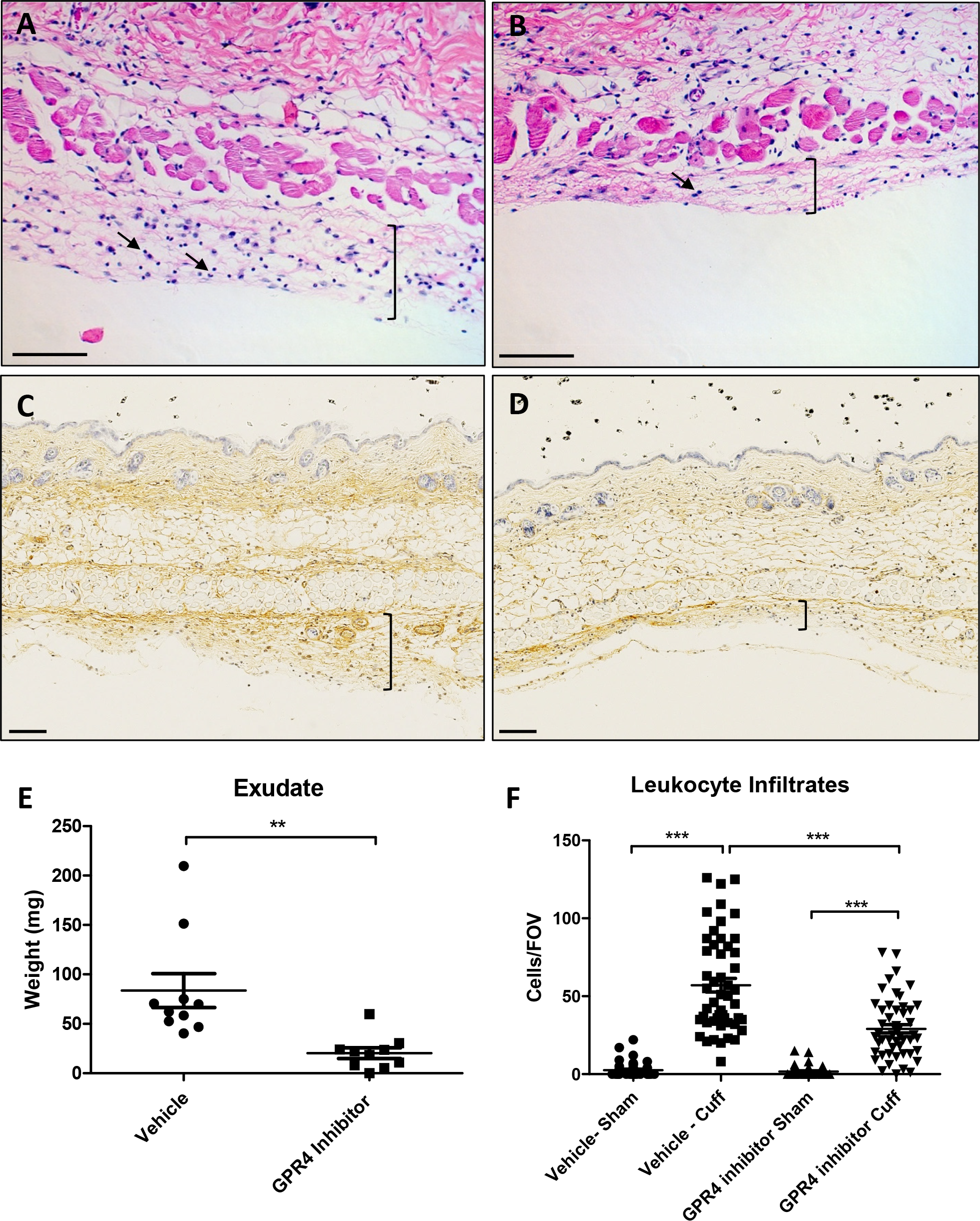
GPR4 antagonist 13 reduces inflammatory exudate production and leukocyte infiltration in the ischemic hindlimb tissues. Inflammatory exudate measurements and leukocyte infiltration quantification in WT mice provided with vehicle or GPR4 antagonist 13 in the hindlimb ischemia-reperfusion mouse model. GPR4 antagonist 13 reduces inflammatory exudate formation and leukocyte infiltration in the exudate. Representative images of H&E staining of (A) WT vehicle cuff and (B) WT GPR4 antagonist 13 cuff skin tissue sections. Immunohistochemical staining of mouse plasma IgG protein in (C) WT vehicle cuff and (D) WT GPR4 antagonist 13 cuff inflammatory exudate sections. IgG protein can be visualized as brown signal. (E) total exudate formation and (F) leukocyte infiltration from multiple fields of view (FOV) when compared to vehicle control following ischemia-reperfusion. Black brackets indicate exudate distribution. Arrows indicate infiltrated leukocytes. 10× and 20× microscope objectives used. Scale bars indicate 100µm. Data analyzed from 10 WT and 10 GPR4 KO mice. Data are presented as mean ± SEM and was analyzed for statistical significance using the unpaired two-tailed *t*-test or ANOVA where **p<0.01 and ***p<0.001.

**Figure 10.**
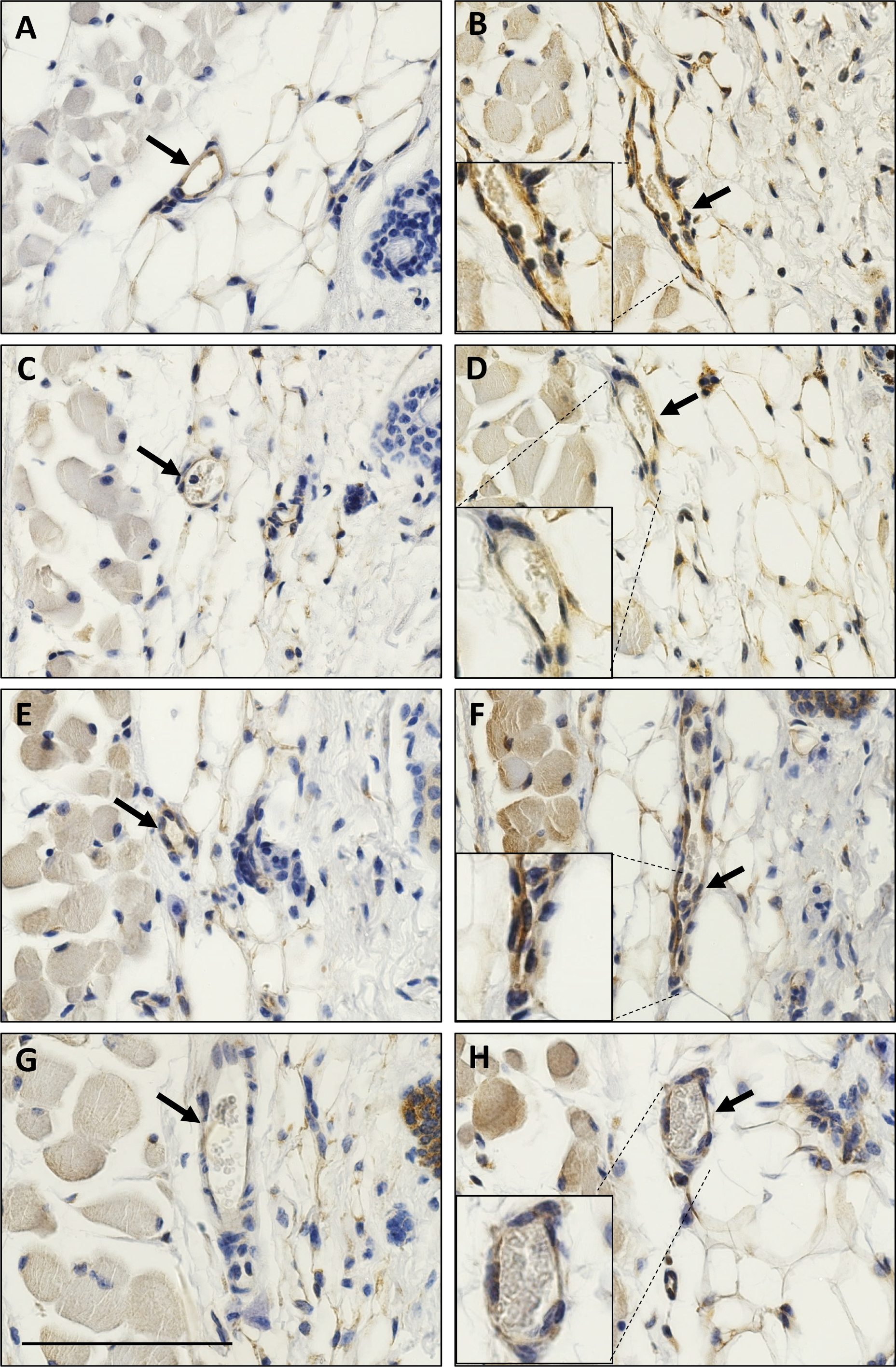
Immunohistochemistry of VCAM-1 and E-selectin protein expression in the loose connective dermal tissue of wild-type mice treated with GPR4 antagonist 13 or vehicle. Immunohistochemical analysis of adhesion molecules VCAM-1 and E-selectin protein expression in vascular endothelial cells within mouse dermal tissue sections of sham and cuff affected hindlimbs. WT mice treated with GPR4 antagonist 13 had reduced expression of VCAM-1 and E-selectin in vascular endothelial cells in cuff-affected tissues when compared to vehicle control. Representative pictures of VCAM-1 signal in (A) WT vehicle sham, (B) WT vehicle cuff, (C) WT GPR4 antagonist 13 sham, and (D) WT GPR4 antagonist 13 cuff dermal connective tissues. Representative pictures of E-selectin expression can be visualized in (E) WT vehicle sham, (F) WT vehicle cuff, (G) WT GPR4 antagonist 13 sham, and (H) WT GPR4 antagonist 13 cuff dermal tissues. Pictures taken with 40× microscope objective. Scale bar indicates 100µm. Arrows indicate blood vessels.

## Discussion

This study demonstrates that activation of GPR4 by acidic pH can stimulate endothelial inflammation through the mediation of paracellular gap formation and permeability by the Gα_12/13_ pathway. We next evaluate the functional role of GPR4 in the inflammatory response within the hindlimb ischemia-reperfusion mouse model using GPR4-null mice and a selective GPR4 antagonist [25]. These data suggest that both the genetic knockout and pharmacologic inhibition of GPR4 reduce the inflammatory response in the ischemia-reperfusion mouse model. These results indicate GPR4 could be a valuable therapeutic target for the remediation of inflammatory disease states.

The inflamed tissue microenvironment has been characterized by a loss of pH homeostasis. Acidosis is a microenvironmental stress factor in which both stromal and infiltrated immune cells exist and by which alters cellular function [7, 12, 16]. Interestingly, local acidosis has been described as a danger signal within ischemic and inflamed tissues which can promote inflammation [36]. However, the way by which cells sense the altered acidic microenvironment and subsequently alter the inflammatory response has only recently been investigated. Cells can employ distinct acid sensing mechanisms such as transient receptor potentials (TRPs), acid sensing ion channels (ASICs) and proton-sensing G protein-coupled receptors (GPCRs) [7, 15, 16, 37]. Proton-sensing GPCRs are activated by acidic pH through the protonation of several histidine residues on the extracellular domains of the receptors which can induce conformational changes in the GPCRs for subsequent G protein activation and downstream signaling [18, 38]. The family of proton-sensing GPCRs include GPR4, GPR65 (TDAG8), and GPR68 (OGR1). GPR65 and GPR68 are predominately expressed in immune cells while GPR4 is expressed in endothelial cells [7, 11, 16, 26]. This family of receptors have only recently been implicated in the regulation of post-ischemic cellular responses and inflammation [16, 39–42].

For example, GPR68 has been suggested to have a cardioprotective function by establishing a cellular protective border zone around the infarcted tissues [42]. GPR65 expression was increased in rat brains and inhibited neuronal apoptosis and post-ischemic inflammatory responses following the middle cerebral artery occlusion and reperfusion event [40]. Studies investigating the role of GPR4 under ischemic conditions are predominately focused on the kidney [39]. A recent study demonstrated that GPR4 deficiency improved renal ischemia-reperfusion injury clinical parameters such as survival rate, serum creatinine, and blood urea nitrogen levels. Furthermore, genetic deficiency of GPR4 deterred apoptosis by suppressing CHOP expression in the kidney [21]. This study is consistent with our previous report demonstrating CHOP expression is dependent on GPR4 activation in HUVECs [5, 6]. However, these studies did not investigate the role of GPR4 in post-ischemic vessel permeability or inflammation.

Our current study reveals a novel role for GPR4 in governing endothelial paracellular gap formation and permeability in response to acidic stress through the Gα_12/13_ pathway. Using both genetic and pharmacological approaches, we demonstrated acidosis-induced GPR4 activation significantly increases the paracellular gap formation and permeability when compared to physiological pH *in vitro*. A previous study reported that GPR4 can reduce endothelial cell barrier function via lysophosphatidylcholine (LPC) [43]. However, the LPC-GPR4 ligand binding interaction were unable to be confirmed [44]. Instead, GPR4 has been shown to function as a pH sensor [17, 18]. Altered endothelial cells permeability can lead to detrimental complications, such as tissue edema. Many diseases related to ischemia-induced inflammation are associated with increased vascular permeability such as stroke, myocardial infarction, sepsis, and cancer [45]. Our study suggests that GPR4 mediates endothelial paracellular gap formation and permeability in response to acidotic stress within the acidic inflamed microenvironment. For this reason, we further investigated the functional role of GPR4 in mediating acute inflammation using the hindlimb ischemia-reperfusion mouse model.

In order to investigate GPR4 in the ischemia-reperfusion model, we used the GPR4 knockout (KO) mice for comparison to wild-type (WT) mice. Our data demonstrated that GPR4 KO mice had reduced parameters of acute inflammation such as tissue edema, inflammatory exudate formation, leukocyte infiltration, and EC adhesion molecule expression (VCAM-1 and E-selectin). These results suggest GPR4 mediates ischemia/reperfusion-induced inflammation most likely through vascular inflammatory programs such as increased vessel permeability and adhesion molecule expression. We and others have previously demonstrated that GPR4 regulates endothelial cell inflammation and ER stress responses [4–6, 8]. Activation of GPR4 in human endothelial cells by acidic pH increased numerous inflammatory cytokines, chemokines, cellular adhesion molecules, and ER stress related genes [5, 6]. Additionally, activation of GPR4 in HUVECs functionally mediated EC-leukocyte interactions of which is necessary for the leukocyte extravasation process in the inflammatory response [4, 6]. The role of GPR4 in the regulation of inflammation has recently been evaluated *in vivo* using the dextran sulfate sodium (DSS)-induced colitis mouse model [19, 20]. GPR4 knockout mice were protected from intestinal inflammation and had reduced adhesion molecule expression in intestinal microvascular endothelial cells when compared to wild-type mice [19]. These data suggest GPR4 potentiates inflammation likely through increased vascular endothelial cell inflammatory responses and are consistent with observations made in this current study and implicate GPR4 as a potential therapeutic target for the remediation of acute and chronic inflammation.

Previous reports have indicated a group of imidazopyridine derivatives, found to selectively inhibit GPR4, can reduce both endothelial cell inflammation *in vitro* and tissue inflammation *in vivo* [5, 6, 8, 46, 47]. We previously demonstrated that GPR4 inhibitors can inhibit GPR4 activation in HUVECs following acidotic stimulation *in vitro* which resulted in reduced expression of GPR4 mediated proinflammatory cytokines, chemokines, and cellular adhesion molecules [5]. Previous *in vivo* studies evaluated GPR4 inhibitors in myocardial infarction, arthritis, nociception, and angiogenesis mouse models and demonstrated that GPR4 inhibition reduced the disease severity when compared to vehicle control [46, 47]. Recently, GPR4 antagonist 13, a pyrazolopyrimidine derivative was developed by Novartis Pharmaceuticals as the next generation of GPR4 inhibitors and found to be more selective for GPR4 inhibition and orally active [25]. GPR4 antagonist 13 was also tested against other pH-sensing GPCRs, the H3 receptor, and hERG channel and demonstrated high selectivity for GPR4. The *in vivo* pharmacokinetics were also evaluated for GPR4 antagonist 13 and found to have good profiles of oral delivery and clearance. GPR4 antagonist 13 was found to effectively reduce arthritic inflammation, hyperplasia, angiogenesis, and colitis [25, 48].

We sought to evaluate the anti-inflammatory effects of GPR4 antagonist 13 within the hindlimb ischemia-reperfusion mouse model. Notably, we observed similar effects of the pharmacological inhibition of GPR4 with GPR4 antagonist 13 when compared to the genetic knockout of GPR4. Our results demonstrated that GPR4 exacerbated post-ischemia/reperfusion tissue inflammation. GPR4 antagonist 13 administration resulted in a decrease in gross edema clinical parameters, inflammatory exudate formation, and leukocyte infiltration. Moreover, GPR4 antagonist 13 treatment reduced endothelial permeability as evidenced by a decrease in plasma IgG protein leakiness when compared to vehicle control. Immunohistochemical analysis of proinflammatory modulators such as endothelial adhesion molecules (VCAM-1 and E-selectin) were also decreased by GPR4 antagonist 13 compared to vehicle.

Taken together, this study demonstrates that GPR4 activation can induce endothelial paracellular gap formation and permeability *in vitro*. Furthermore, evaluation of the contribution of GPR4 in the hindlimb post-ischemia/reperfusion inflammatory response supports the role for GPR4 in vascular permeability and inflammation. Genetic knockout and pharmacological inhibition of GPR4 *in vivo* can decrease leukocyte infiltration and the expression of endothelial adhesion molecule VCAM-1 and E-selectin, and reduce vascular permeability as evidenced by attenuated plasma IgG leakiness into the subcutaneous connective tissues and exudate formation. The results suggest that inhibition of GPR4 can be exploited as a novel approach to alleviate inflammation and tissue edema.

## Acknowledgement

This study was supported in part by research grants from the National Institutes of Health (R15DK109484, to L.V.Y.) and the American Heart Association (11SDG5390021, to L.V.Y.). We thank Nancy Leffler, Lixue Dong, Joani Oswald, Comparative Medicine staff, Drs. Yan-Hua Chen, Warren Knudson, David Tulis, Karen Oppelt, and Kvin Lertpiriyapong for helpful discussion and technical assistance. We also thank the Novartis Institutes for BioMedical Research for providing the GPR4 antagonist 13 and Dr. Owen Witte for the GPR4 knockout mice.

## Author contributions

E.A.K. and L.V.Y. designed the experiments; E.A.K., E.J.S. and M.A.M. performed the experiments; E.A.K., E.J.S., M.A.M. and L.V.Y. analyzed the data; L.V.Y. supervised the research; J.V. and P.L. provided the GPR4 antagonist 13 and instructions on how to administer the compound in mice; E.A.K., E.J.S. and L.V.Y. wrote the manuscript; all authors reviewed the manuscript.

---

**Supplemental Figure 1.**
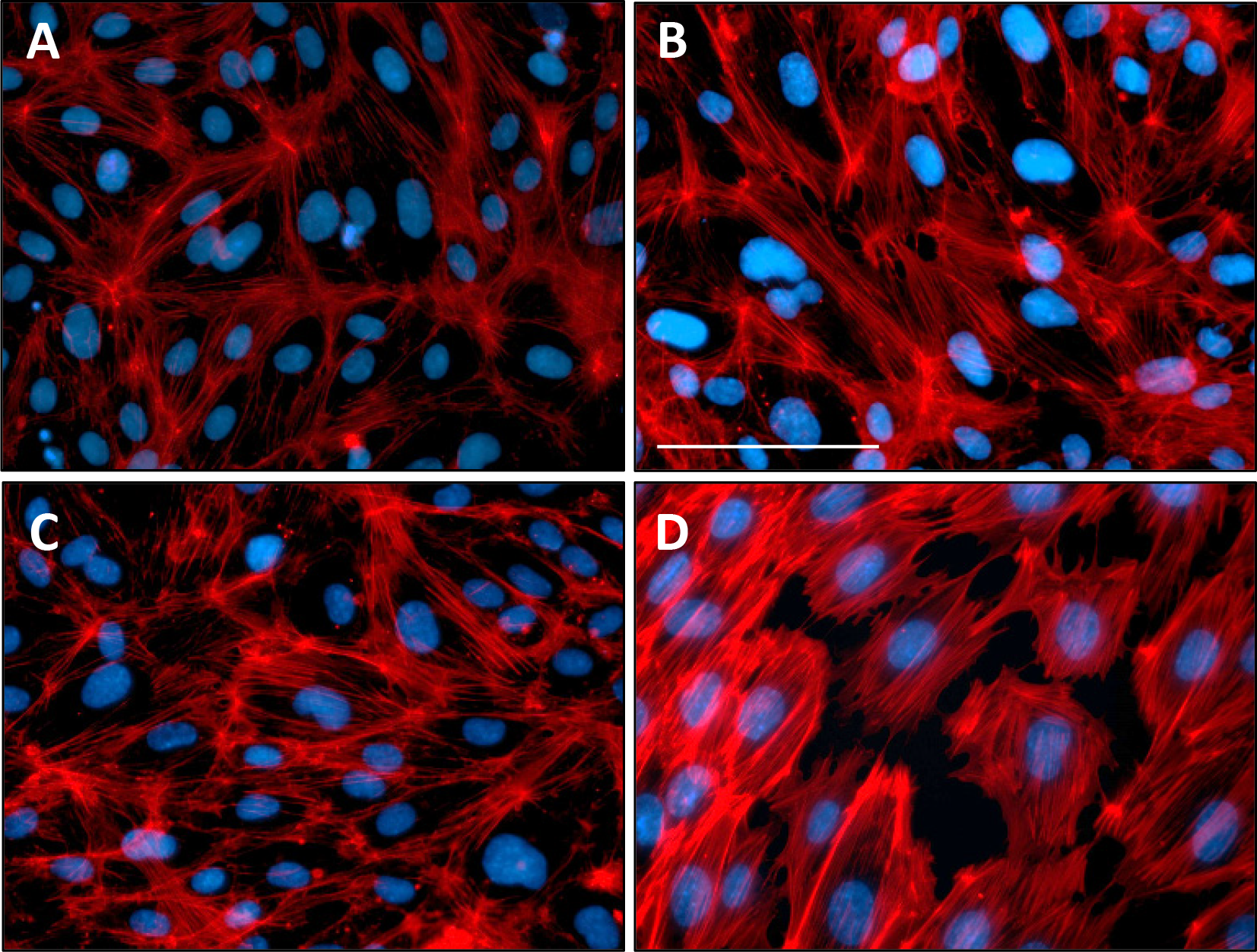
GPR4 activation by acidic pH induces actin stress fiber formation in HUVECs. Actin stress fiber formation in HUVECs treated with physiological or acidic pH. Activation of GPR4 by acidic pH 6.4 increases actin stress fibers when compared to physiological pH 7.4 in HUVECs. Representative images of F-actin labeled with rhodamine phalloidin in (A) HUVEC/vector pH 7.4 or (B) pH 6.4 and (C) HUVEC/GPR4 pH 7.4 or (D) pH 6.4. HUVECs were treated with pH stimulation medium for 5 hours. Scale bar indicates 100µm.

**Supplemental Figure 2.**
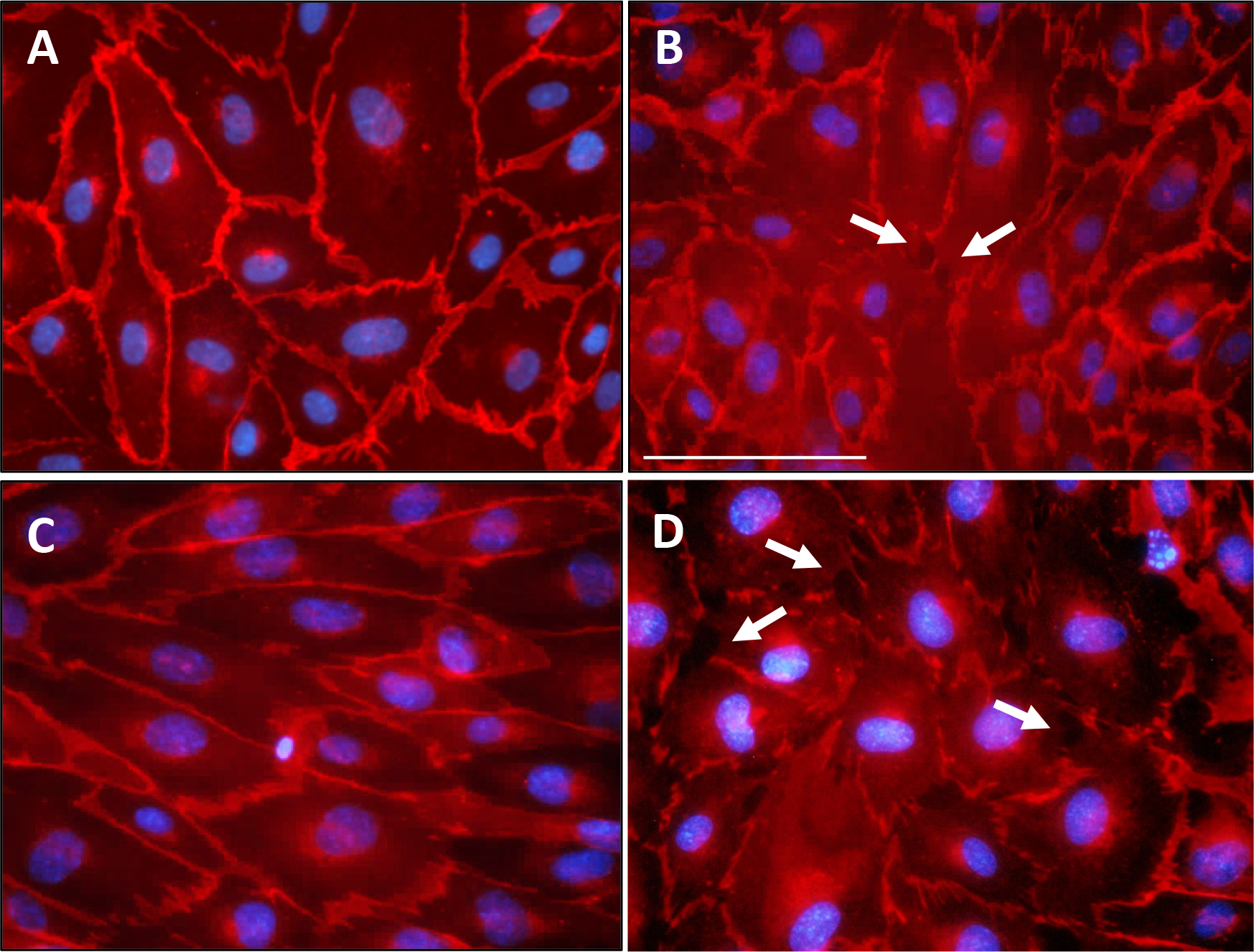
VE-Cadherin protein expression is reduced on endothelial paracellular gap borders. VE-Cadherin protein expression in HUVECs treated with physiological or acidic pH. Acidosis/GPR4-mediated endothelial paracellular gap areas have reduced VE-Cadherin expression along endothelial cell borders. Representative images of VE-Cadherin expression in (A) HUVEC/vector pH 7.4 or (B) pH 6.4 and (C) HUVEC/GPR4 pH 7.4 or (D) pH 6.4. HUVECs were treated with pH stimulation medium for 5 hours. Scale bar indicates 100µm. Arrows indicate endothelial cell borders with reduced VE-Cadherin protein expression.

**Supplemental Figure 3.**
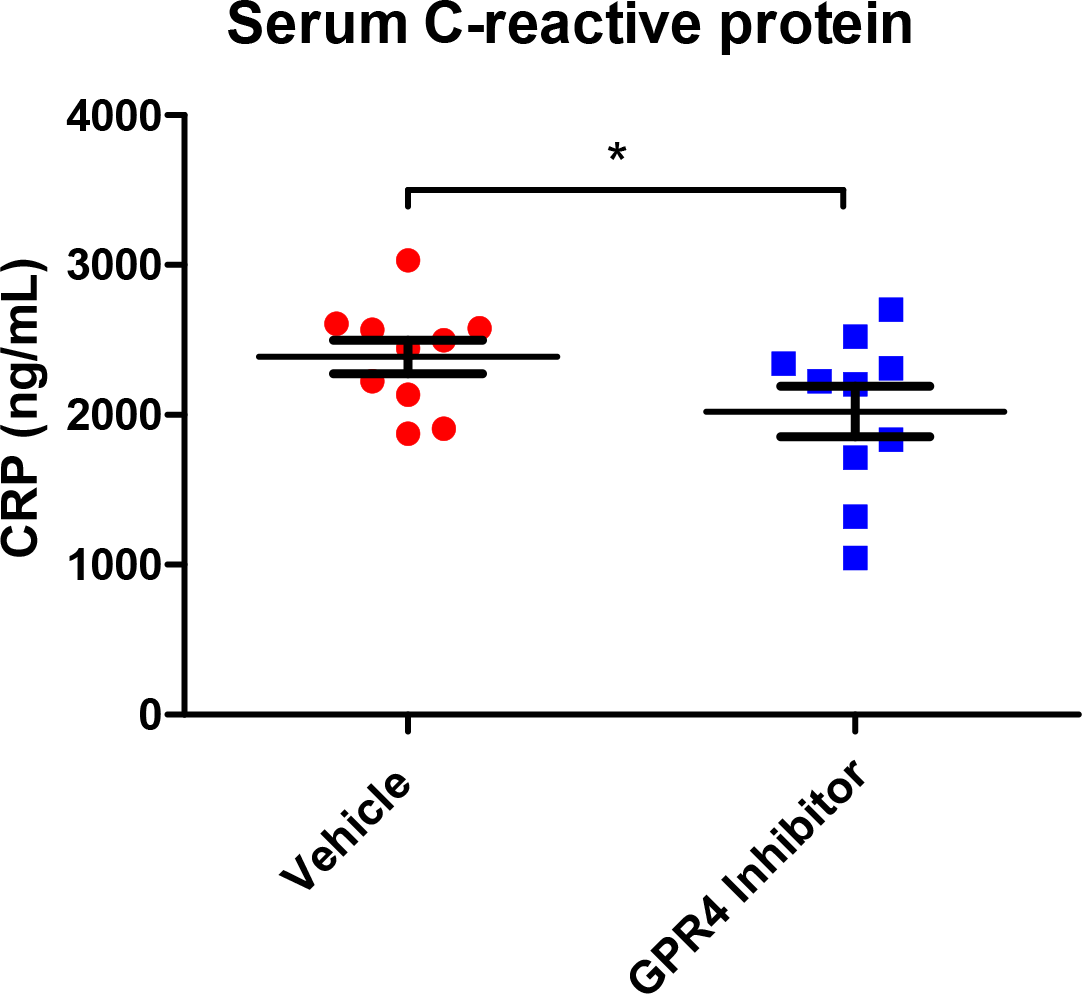
Mice treated with GPR4 antagonist 13 have reduced serum C-reactive protein levels when compared to vehicle control. C-reactive protein (CRP) inflammatory marker in mouse serum. As previous indicators suggest GPR4 antagonist 13 reduces inflammation in the hindlimb ischemia-reperfusion mouse model, we assessed the anti-inflammatory effects of GPR4 antagonist 13 in the reduction of CRP levels in mouse serum. Mouse serum CRP was measured by ELISA and represented as ng/mL. Data analyzed from 10 WT vehicle and 10 WT GPR4 antagonist 13 mice. Data are presented as mean ± SEM and analyzed for statistical significance using the unpaired one-tailed *t*-test where *p<0.05.

